# Evasion of CARD8 Activation During HIV-1 Assembly

**DOI:** 10.1101/2025.05.19.654850

**Authors:** Ivy K. Hughes, James B. Hood, Andrés A. Quiñones-Molina, Hisashi Akiyama, Suryaram Gummuluru

## Abstract

As intracellular parasites, viruses must devise sophisticated mechanisms to produce and assemble viral components while suppressing activation of innate immune effectors. Here, we report that coordination of HIV-1 assembly by the viral polyprotein Gag suppresses inappropriately-timed protease (PR) activity to evade the PR activity sensor, CARD8. Employing mutants of Gag, we show that disruption of domains controlling viral assembly site (MA) or virus particle release (NC and p6) lead to premature activation of PR and the CARD8 inflammasome, resulting in IL-1β secretion and pyroptotic cell death. Further, we demonstrate that previously-observed host-adaptive mutations in HIV-1 MA (M30K) and p6 (PTAP duplication) associated with greater fitness in humans differentially modulate the process of viral assembly and budding to evade CARD8-mediated cell death. Altogether, this work reveals adaptation to human CARD8 by HIV-1 Gag upon zoonotic transmission from chimpanzees and suggests that assembly-regulated CARD8 activation influences the trajectory of HIV-1 evolution and fitness in humans.

## INTRODUCTION

Human immunodeficiency virus type 1 (HIV-1) encounters multiple innate immune sensors during viral replication, recognition by which can lead to the activation of type I interferon (IFN) and pro-inflammatory signaling cascades and cytokine secretion. Activation of innate immune pathways is most often deleterious to viral replication, particularly during early events of viral transmission at mucosal surfaces.(*1*) Multiple cell-intrinsic antiviral IFN-stimulated genes (ISGs) have been shown to restrict productive HIV-1 infection, including antagonism of viral budding by tetherin (counteracted by HIV-1 Vpu) and targeted hypermutation of viral genomes by APOBEC3G (counteracted by HIV-1 Vif), highlighting the critical importance of IFN resistance displayed by successful transmitter/founder strains of HIV-1.(*2*) In addition to IFN-inducing sensors, cell-intrinsic inflammasome sensors also restrict virus infection via induction of pyroptosis of productively infected cells, which eliminates virus-producing cells and results in the secretion of pro-inflammatory cytokines IL-1β and IL-18.(*3*)

Among these cell-intrinsic inflammasome sensors is CARD8, which mimics HIV-1 protease (PR) cleavage site specificity and initiates pyroptosis in response to intracellular viral proteolytic activity.(*4*) CARD8 patrols the cytoplasm as a non-covalent complex of an N-terminal fragment (CARD8-N) and a C-terminal fragment (CARD8-C). Upon proteolysis of the N-terminal fragment by viral proteases, including HIV-1 PR, the C-terminal fragment is liberated and forms the CARD8 inflammasome which initiates pyroptosis.(*5*) The detection of HIV-1 PR activity by CARD8 was first reported in the context of the treatment of productively infected cells with non-nucleoside reverse transcription inhibitors (NNRTIs).(*4*) NNRTIs artificially enforce Gag-Pol dimerization and inappropriate activation of cytoplasmic PR activity.(*6*) Activation of CARD8 by incoming virion-associated HIV-1 PR has also been reported, and is initiated upon virus particle fusion with target cells, particularly at high multiplicities of infection (MOIs) or during cell-to-cell transmission.(*7*) However, the activation of CARD8 or lack thereof during the process of viral particle assembly has until this point not been investigated.

HIV-1 assembly begins with recognition of the phosphatidyl inositol (4,5) bisphosphate (PI(4,5)P_2_)-rich plasma membrane (PM) by the matrix (MA) domain of Gag.(*8*) Additional Gag and Gag-Pol monomers as well as unspliced genomic RNA are recruited to the assembly site, and capsid (CA)- and nucleocapsid (NC)-mediated interactions drive Gag oligomerization and particle assembly.(*9*) Recruitment of ESCRT factors Tsg101 and/or ALIX by p6 results in timely budding, completing the assembly process.(*10*) PR is required for final maturation of the viral particle into an infectious state.(*11*) The timing of HIV-1 PR activity is tightly controlled, and the dimeric PR does not activate until the moment of or slightly after viral particle release.(*12, 13*) This tight control over the timing of PR activity, combined with the apparent lack of CARD8 activation in the absence of NNRTI treatment, together strongly suggest the existence of virus-encoded mechanisms to restrain PR activity to avoid spurious CARD8 activation during viral assembly. To investigate these mechanisms of virus-encoded control of PR activity, we performed *in vitro* studies using cell lines and primary cells infected with viruses encoding mutants of Gag. Strikingly, we found that multiple regions of Gag, namely MA, NC, and p6, regulate viral proteolytic activity and CARD8 activation. Remarkably, we also find that previously documented mutations in the MA and p6 regions of Gag associated with greater replicative fitness in humans improve evasion of pyroptotic cell death in HIV-infected cells. In summary, this work reveals a previously undescribed evasion mechanism of HIV-1 from the cell-intrinsic innate immune system and highlights novel domains of Gag potentially amenable to pharmaceutical intervention to kill infected cells.

## RESULTS

### HIV-1 viral matrix protein suppresses inflammasome activation

HIV-1 matrix (MA) is composed of a lipidated N-terminal myristoyl moiety followed by a series of basic residues known as the highly-basic region (HBR). HBR displays a specific recognition capacity for the PI(4,5)P_2_-rich inner leaflet of the plasma membrane (PM). In addition, HBR coordinates temporal binding of cellular tRNAs to suppress promiscuous binding of Gag to internal membranes and spontaneous myristoyl exposure, thus allowing for specific PM recognition.(*14, 15*) As the driver of membrane recognition and initial steps of membrane-associated virus assembly, we chose to first analyze the role of MA in CARD8 activation. We began our investigation by overexpressing caspase-1, pro-IL-1β, and single-round HIV-1 proviral constructs in HEK293T cells by transient transfection. We analyzed inflammasome activation by measuring pro-IL-1β cleavage and IL-1β secretion by Western blot and ELISA, respectively. Overexpression of wild-type (WT) HIV-1 in HEK293T cells, which leads to spontaneous dimerization of Gag-Pol, resulted in intracellular pro-IL-1β processing and IL-1β secretion (**Fig. 1A/B**). Interestingly, overexpression of a Rev-inactive virus (M10) which attenuates cytoplasmic Gag/Gag-Pol expression (**Fig. 1A** and (*16*)) or a myristoylation-deficient MA (G2A) virus mutant which attenuates membrane-association of Gag,(*16*) abrogated IL-1β secretion (**Fig. 1A**), consistent with a model where CARD8 is activated by viral PR generated during the process of *de novo* membrane-associated virus assembly under overexpression conditions. To analyze MA’s role in the regulation of viral PR activity during assembly, we employed a MA mutant where all amino acids except the myristoylation signal and MA-CA PR cleavage site have been removed (ΔMA6-125).(*16, 17*) Strikingly, overexpression of ΔMA6-125 resulted in a significant increase in pro-IL-1β processing (**Fig. 1C**) and IL-1β secretion (**Fig. 1D**) compared to WT. This effect was dependent on viral PR, as catalytically-inactive mutant of PR combined with the ΔMA6-125 mutation (ΔMA6-125pro-) showed a lack of Gag processing, pro-IL-1β cleavage, and IL-1β secretion (**Fig. 1C/D**).

**Fig. 1:**
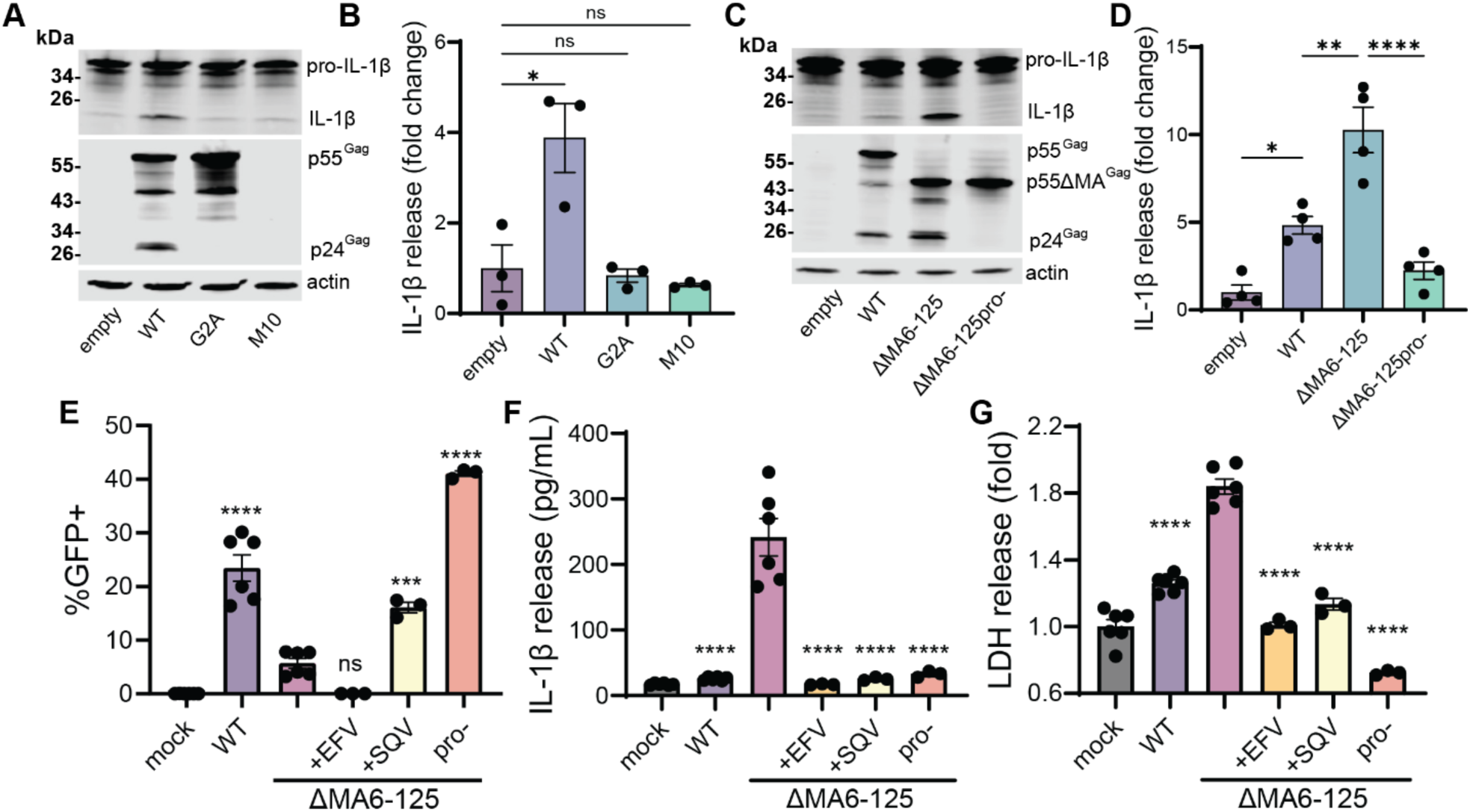
Matrix-deficient HIV-1 assembly induces inflammasome activation. (**A**-**D**) HEK293T cells were transfected with plasmids encoding caspase-1, pro-IL-1β, and the indicated single-round proviral constructs. Western blots show pro-IL-1β cleavage in cell lysates with HIV-1 protein expression assayed with anti-p24^Gag^ antibody (**A**/**C**). IL-1β secretion in cell supernatants (**B**/**D**) was determined by ELISA at 24h post-transfection. (**E**-**G**) PMA/THP macrophages were co-infected with VSV-G-pseudotyped single-round HIV-1Δenv/GFP and Vpx-containing SIV3+ VLPs. Cells were stimulated with TNFα after infection and plots show GFP positivity assayed by flow cytometry (**E**), or IL-1β (**F**) and LDH (**G**) secretion at 3 days post-infection. +EFV/SQV refers to ΔMA6-125 infection in the presence of the indicated antiretroviral, and pro-refers to ΔMA6-125pro-virus infection. Data is shown as mean +/- SEM. One-way ANOVA was used with Tukey’s post-test (**B**/**D**) or Dunnett’s post-test (**E**-**G**): ns: p≥0.05; *: p<0.05; **: p<0.01; ***: p<0.001; ****: p<0.0001. In **E**-**G**, statistics refer to comparison with ΔMA6-125 infection. (**A**-**D**) Western blots and cytokine release are from a single experiment which is representative of three independent experiments. (**E**-**G**) Data points are individual wells across two experiments, representative of over three independent experiments.

We then sought to confirm this effect in a model of HIV-1 infection. To this end, we differentiated monocytic THP-1 cells into macrophage-like cells by treatment with PMA (PMA/THP). We then co-infected PMA/THP cells with single round (Env-deficient), GFP-expressing, VSV-G pseudotyped HIV-1 and SIVmac Vpx-containing virus-like particles (SIV3+ VLPs, to overcome SAMHD1-mediated restriction of HIV-1 in myeloid cells).(*18*) To normalize early post-entry behavior between WT and mutant viruses, all mutant viruses were produced with WT HIV-1 Gag/Pol packaging plasmid. Three days post-infection, we observed a notable failure in survival of infected GFP+ cells in ΔMA6-125-infected cultures which was rescued upon treatment with HIV-1 protease inhibitor saquinavir (SQV) or PR inactivation (**Fig. 1E**). Further, cells infected with ΔMA6-125 virus secreted high levels of IL-1β, which was not observed in WT-infected PMA/THP cells (**Fig. 1F**). IL-1β secretion in ΔMA6-125 virus infected-PMA/THP macrophages required HIV-1 PR activity as it was completely abolished by SQV treatment or catalytic inactivation of PR (ΔMA6-125pro-). Inflammasome activation was also not dependent on the incoming ΔMA6-125 virus particle-associated PR, as IL-1β secretion was completely abrogated by treatment with HIV-1 reverse transcriptase (RT) inhibitor efavirenz (EFV) prior to the initiation of infection (**Fig. 1F**), which does not block entry of virus particles or the deposition of viral PR into the cytoplasm. Further, compared to WT-infected cells, ΔMA6-125-infected cells secreted higher levels of lactate dehydrogenase (LDH, **Fig. 1G**), indicating pyroptosis of infected cells and extracellular release of cytoplasmic contents. These experiments indicate that MA plays an important role in restraining PR activity and evasion from inflammasome activation during HIV-1 infection of macrophages.

### Assembly-driven inflammasome activation is orchestrated by CARD8

The dependence on PR for inflammasome activation in ΔMA6-125-infected cells suggested that PR-sensing CARD8 was the inflammasome sensor responsible for the phenotype. To confirm this, we repeated overexpression experiments in HEK293T-CARD8KO cells (with the CARD8 open reading frame disrupted via CRISPR) and infection experiments with PMA/THP-shCARD8 cells (where CARD8 expression was knocked down via transduction with CARD8shRNA-expressing lentivector) (**Fig. 2A**). Strikingly, in comparison to parental HEK293T cells, pro-IL-1β cleavage and IL-1β secretion were abrogated in ΔMA6-125-transfected HEK293T-CARD8KO cells (**Fig. 2B/C**). Similarly, in PMA/THP-shCARD8 cells, IL-1β (**Fig. 2E**) and LDH secretion (**Fig. 2F**) were abolished and survival of GFP+ cells (**Fig. 2D**) was similar between the WT and ΔMA6-125 virus-infected cultures. These results confirm that CARD8 is the sensor activated during MA-deficient virus assembly in HEK293T cells and PMA/THP macrophages.

**Fig. 2:**
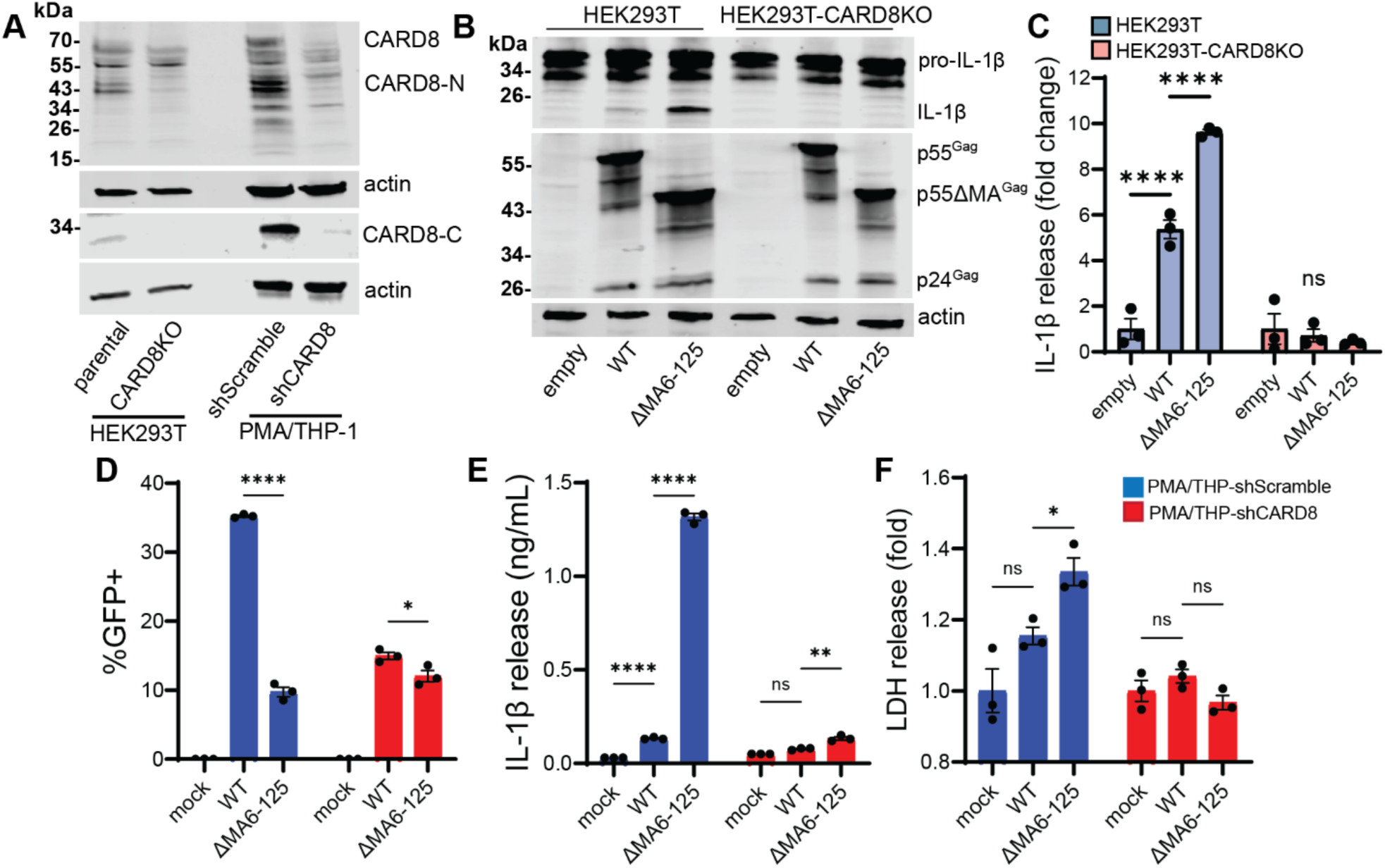
CARD8 activation during matrix-deficient HIV-1 assembly. (**A**) Western blots showing HEK293T or PMA/THP cell lysates probed with anti-CARD8-N or anti-CARD8-C antibodies. (**B**) Western blots showing pro-IL-1β cleavage in transfected HEK293T cells and HIV-1 protein expression assayed with anti-p24^Gag^ antibody 24h post-transfection of single-round proviral plasmid, caspase-1, and pro-IL-1β. (**C**) IL-1β secretion from HEK293T or HEK293T-CARD8KO cells at 24h post-transfection. (**D**-**F**) GFP positivity, IL-1β secretion, and LDH secretion at 3 days post-infection of TNFα-treated PMA/THP cells stably transduced with shScramble- or shCARD8-encoding lentivector and co-infected with single-round VSV-G-pseudotyped HIV-1Δenv/GFP and SIV3+ VLPs. Data is shown as mean +/- SEM. Two-way ANOVA was used with Tukey’s post-test: ns: p≥0.05; *: p<0.05; **: p<0.01; ****: p<0.0001. (**B**/**C**) Western blots and cytokine release are from a single experiment which is representative of three independent experiments. (**D**-**F**) Data points are individual wells from one experiment, which is representative of three independent experiments.

### Viral assembly is required for CARD8 activation in MA-deficient virus infection

PR activation occurs during the process of viral assembly and maturation, when Gag-Pol monomers are recruited to the membrane-associated virus assembly site and PR domains of adjacent Gag-Pol molecules come in close proximity to dimerize and become enzymatically active. Hence, we sought to determine whether initiation of virus assembly was needed for CARD8 activation. To this end, we added additional mutations in the context of ΔMA6-125 that disrupt early steps of virus assembly: the myristoylation-deficient mutant (G2A) that prevents membrane association of Gag, and a C-terminal capsid (CA-CTD) mutation (VK181/2AA) that inactivates CA-CA dimerization needed for the formation of higher-order CA oligomers. (*16*) We found that these mutations significantly reduced pro-IL-1β cleavage and IL-1β secretion in ΔMA6-125 overexpressing HEK293T cells (**Fig. 3A**-**D**) and significantly reduced IL-1β and LDH secretion in ΔMA6-125-infected PMA/THP macrophages (**Fig. 3E**-**G**), suggesting that the source of the CARD8-activating viral PR in infected cells was indeed the membrane-associated virus assembly site. To determine whether CARD8 activation strictly required membrane-associated PR activity, we induced CARD8 activation via NNRTI (rilpivirine, RPV) treatment in HIV-1/WT or G2A-infected PMA/THP cells. Surprisingly, we observed similar levels of IL-1β secretion and LDH release in both virus infections (**Fig. S1**), suggesting that enzymatically active membrane-associated PR or cytosolic PR (upon NNRTI-induced activation) can be sensed by CARD8.

**Fig. 3:**
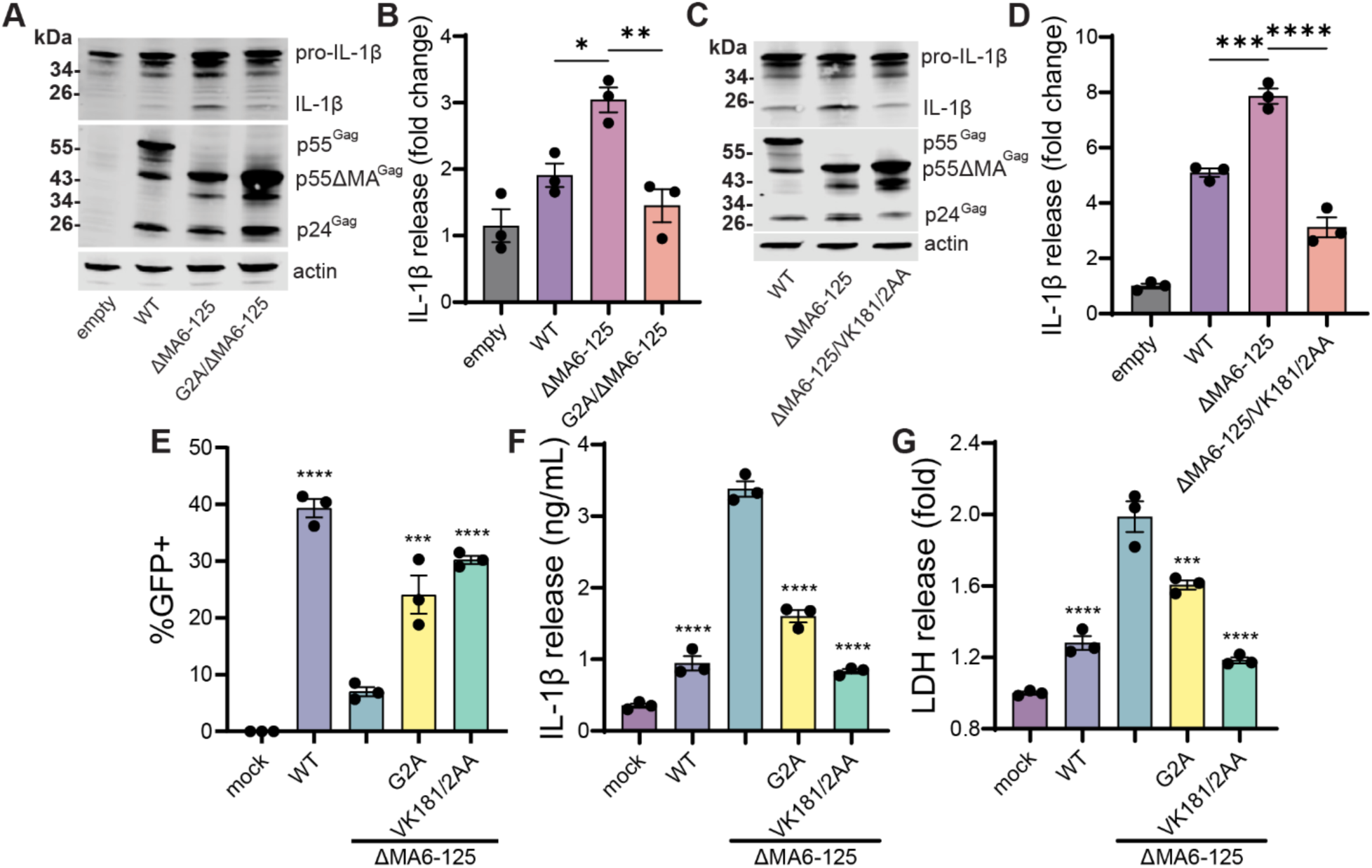
Membrane-associated viral assembly is required for CARD8 activation in MA-deficient virus infection. (**A**/**C**) Western blots showing pro-IL-1β cleavage and HIV-1 protein expression assayed with anti-p24^Gag^ antibody 24h post-transfection of HEK293T cells with single-round proviral plasmid, caspase-1, and pro-IL-1β, with plots (**B**/**D**) showing IL-1β secretion. (**E**-**G**) GFP positivity, IL-1β secretion, and LDH secretion at 3 days post-infection of TNFα-treated PMA/THP cells co-infected with single-round VSV-G-pseudotyped HIV-1Δenv/GFP and SIV3+ VLPs. Data is shown as mean +/- SEM analyzed by one-way ANOVA with Tukey’s post-test (**B**/**D**) or Dunnett’s post-test (**E**-**G**): *: p<0.05; **: p<0.01; ***: p<0.001; ****: p<0.0001. (**A**-**D**) Western blots and cytokine release are from a single experiment which is representative of three independent experiments. (**E**-**G**) Data points are individual wells from one experiment, which is representative of three independent experiments. Statistics refer to comparison with ΔMA6-125 infection.

### p6 modulates CARD8 sensing by preventing PR assembly site escape

Besides MA, nucleocapsid (NC) contributes to immature virus capsid assembly by promoting viral genomic RNA (gRNA) binding-dependent Gag multimerization.(*19*) In addition, basic residues of NC can bind to acidic phospholipids in host membranes and along with p6 can facilitate virus particle release by recruiting cellular ESCRT complexes for membrane scission.(*9, 20*) We reasoned that gRNA recruitment by NC or timely membrane scission driven by NC and p6 could have roles in the initiation or shielding of PR activity, thus preventing CARD8 activation. To inactivate NC, we generated ΔNC15-49 by deleting the entirety of the two zinc finger domains,(*21*) and to inactivate p6 we performed mutagenesis where both Tsg101/ESCRT-I-recruiting PTAP motifs in Lai-p6 were changed to the inert sequence LIRL (named ΔPTAP, **Fig. 4A**).(*22*) We also probed NC function with a mutant in which the entirety of NC sequence was replaced with a nonspecific RNA-interacting leucine zipper domain of a eukaryotic transcription factor (GagZip).(*23*) In PMA/THP macrophages, infection with ΔPTAP mutant virus resulted in dramatically enhanced IL-1β and LDH release, similar to that observed with ΔMA6-125 mutant virus infection (**Fig. 4C/D**). In contrast, inflammasome activation was not observed in ΔNC15-49 or GagZip mutant virus infections (**Fig. 4C/D**). Further, knockdown of CARD8 abrogated IL-1β and LDH secretion in ΔPTAP virus-infected cells (**Fig. 4F/G**). Finally, we observed striking increases in pro-IL-1β cleavage in HEK293T cells overexpressing ΔPTAP mutant (**Fig. 4H**), that was dependent on PR activity (**Fig. 4I**). In contrast, pro-IL-1β cleavage in HEK293T cells was nearly absent in both ΔNC15-49 and GagZip mutants (**Fig. 4H**).

**Fig. 4:**
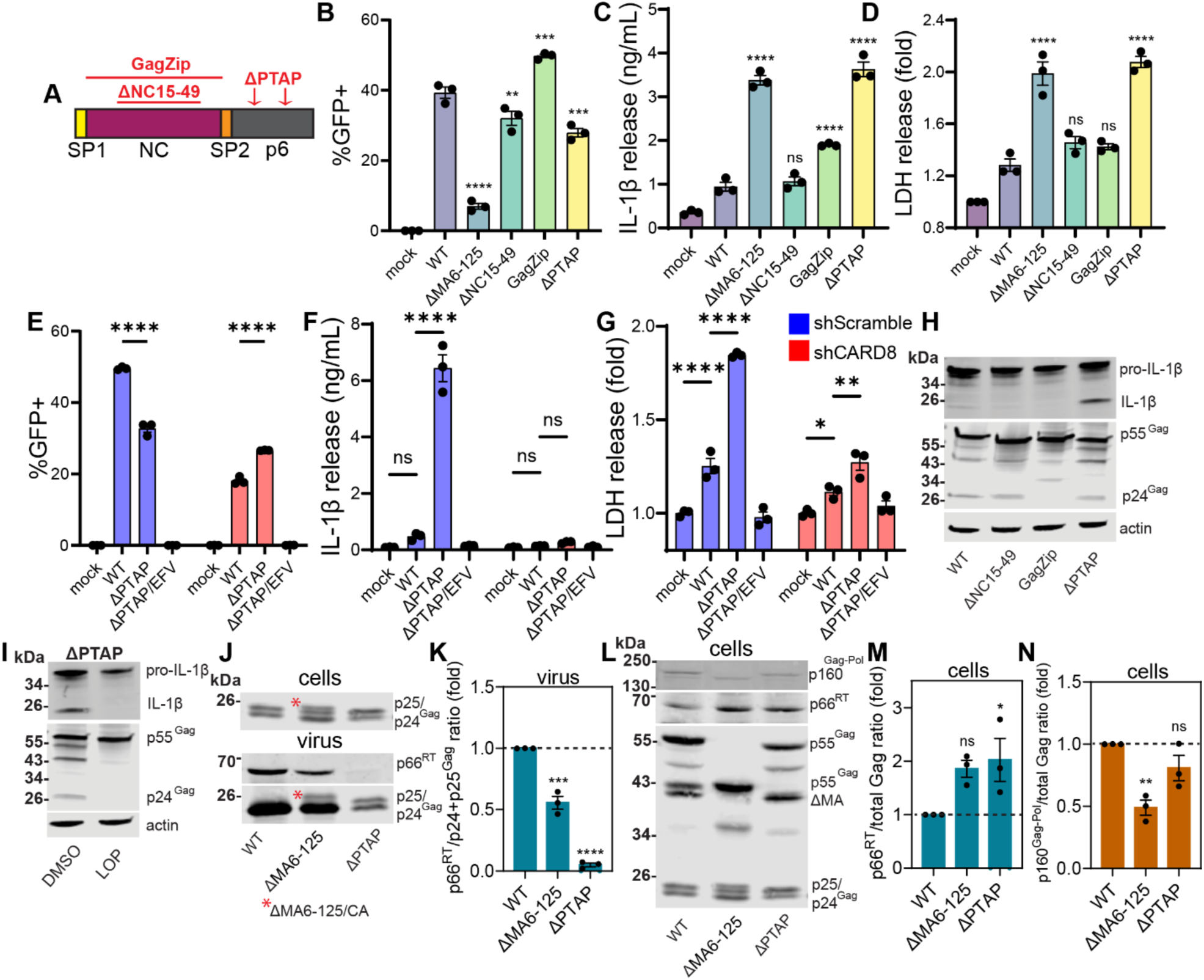
HIV-1 PR escapes the viral assembly site and activates CARD8 during p6-deficient virus assembly. (**A**) Schematic of the C-terminal portion of Gag outlining the virus mutants used in these experiments. (**B**-**D**) PMA/THP cells were co-infected with VSV-G-pseudotyped single-round HIV-1 and SIV3+ VLPs. Cells were stimulated with TNFα after infection and plots show GFP positivity (**B**), IL-1β secretion (**C**), and LDH secretion (**D**) into the culture medium at 3 days post-infection. (**E**-**G**) GFP positivity, IL-1β secretion, and LDH secretion at 3 days post-infection of TNFα-treated PMA/THP cells stably transduced with shScramble- or shCARD8-encoding lentivector and infected as described above. EFV refers to infection in the presence of efavirenz. (**H**) Western blots showing IL-1β cleavage in HEK293T cells, plus HIV-1 protein expression assayed with anti-p24^Gag^ antibody 24h post-transfection of caspase-1, pro-IL-1β, and the indicated single-round proviral construct. (**I**) Western blots showing IL-1β and Gag protein (assayed with anti-p24^Gag^ antibody) of HEK293T cells transfected with pro-IL-1β/caspase-1 plasmids and ΔPTAP single-round proviral plasmid, treated with DMSO or 1 μM lopinavir (LOP). (**J**/**L**) Western blots with cell or ultracentrifuge-pelleted virus lysates from transfected HEK293T cells (provirus-only transfection) showing p160^Gag-Pol^, p66^RT^, p25/p24^Gag^, and total Gag (probed with anti-p24^Gag^, anti-HIV-Ig, anti-p24^Gag^, and anti-p24^Gag^ antibodies respectively), with the ΔMA6-125/CA artificial cleavage product annotated in (**J**). (**K**/**M**/**N**) Quantification from Western blots showing p66^RT^/p24+p25^Gag^ ratio in virus lysates (**K**), p66^RT^/total Gag ratio in cell lysates (**M**), and p160^Gag-Pol^/total Gag ratio in cell lysates (**N**), normalized to WT. Data is shown as mean +/- SEM analyzed with one-way ANOVA with Dunnett’s post-test (**B**-**D**/**K**/**M**/**N**) or two-way ANOVA with Tukey’s post-test (**E**-**G**): ns: p≥0.05; *: p<0.05; **: p<0.01; ***: p<0.001; ****: p<0.0001. (**B**-**D**) Data points are individual wells from three independent experiments, with statistics referring to comparison with WT. (**E**-**G**) Data points are individual wells from one experiment, which is representative of three independent experiments. (**H**/**I**) Western blots are representative of three independent experiments. (**J**-**N**) Western blots are from one representative experiment with quantification statistics referring to comparison with WT across three individual transfection experiments.

Because p6 is responsible for ESCRT recruitment and virus particle budding is roughly ten-fold slower in the absence of p6,(*24*) we hypothesized that CARD8 was being activated by auto-processed PR that escaped the viral assembly site due to defects in timely budding in ΔPTAP virus-infected cells. Previous findings have suggested that ΔPTAP virus particles contain sub-optimal levels of Pol if PR is intact,(*25*) implying viral enzymatic activities escape from membrane-associated assembly sites to the host cell cytoplasm if budding is slow. We additionally hypothesized that a similar defect in particle release or aberrant initiation of Gag-Pol auto-processing could explain the role of MA in evading CARD8 activation. To address these hypotheses, we analyzed Gag/Gag-Pol processing in transfected HEK293T cells and their virus progeny via Western blot analysis. We first reasoned that analyzing the successful complete processing of p25^Gag^ to p24^Gag^ in virus particles, as one of the final PR cleavage events occurring in virus particle maturation after budding, could be indicative of the quantity of PR being retained in the assembling virus particle as opposed to escaping to the host cell cytoplasm. WT, ΔMA6-125, and ΔPTAP virus-transfected cell lysates all contained a mixture of unprocessed p25^Gag^ and p24^Gag^, as expected in a bulk population under overexpression conditions (**Fig. 4J**). In virus particles, however, ΔPTAP was the only mutant in which a significant portion of p25^Gag^ remained unprocessed, consistent with the hypothesis that active PR (after auto-processing of Gag-Pol) escapes from the assembly site due to delays in virus particle budding during ΔPTAP virus assembly. While we observed an extra band in ΔMA6-125-transfected virus pellets, that band was also present in ΔMA6-125-transfected cell lysates (**Fig. 4J**), which we hypothesize likely reflects a failure to process the ΔMA6-125/CA cleavage site as HIV-1 PR is notably dependent on sequence context for cleavage site recognition.(*26, 27*) When we probed virus pellets with anti-HIV-Ig, however, we noticed a loss of p66^RT^ in both ΔMA6-125 and ΔPTAP virus particles (**Fig. 4J/K**), corroborating our hypothesis that there was poor retention of active viral enzymes in both ΔMA6-125 and ΔPTAP virus assembly. Indeed, in both ΔMA6-125 and ΔPTAP transfections, we observed a concomitant increase of processed p66^RT^ in cell lysates, in contrast to virus pellets (**Fig. 4L/M**). Finally, in ΔMA6-125-transfected cell lysates, we observed a decrease of full-length unprocessed p160^Gag-pol^, indicating aberrant initiation of cytoplasmic processing of Gag-pol (**Fig. 4N**). Taken together, these results indicate that both MA and p6 regulate CARD8 sensing of intracellular PR activity during viral assembly in HEK293T cells and macrophages.

### Gag regulates PR-induced CARD8 activation in primary CD4+ T cells

After determining that MA and p6 orchestrated viral assembly to conceal PR activity in infected myeloid and transfected HEK293T cells, we next determined the effects of Gag assembly mutants on inflammasome activation in primary CD4+ T cells. To this end, we infected activated primary CD4+ T cells with VSV-G-pseudotyped, GFP-expressing single-round viruses, and probed CARD8 activity by monitoring survival of infected CD4+ T cells in the presence or absence of the protease inhibitor lopinavir (LOP). Control experiments showed that with WT infection, LOP treatment did not affect survival of infected GFP+ cells. In contrast, death of infected cells (measured by comparative loss of GFP positivity) could be robustly induced by treating WT-infected cells with RPV, and RPV-mediated cytotoxicity could be prevented by pre-treatment with LOP (**Fig. 5A**). To further confirm the activity of CARD8 in activated primary CD4+ T cells, we also treated activated, uninfected CD4+ T cells with the CARD8 activating ligand Val-boroPro (VbP) and observed LDH release (**Fig. S2**).

**Fig. 5:**
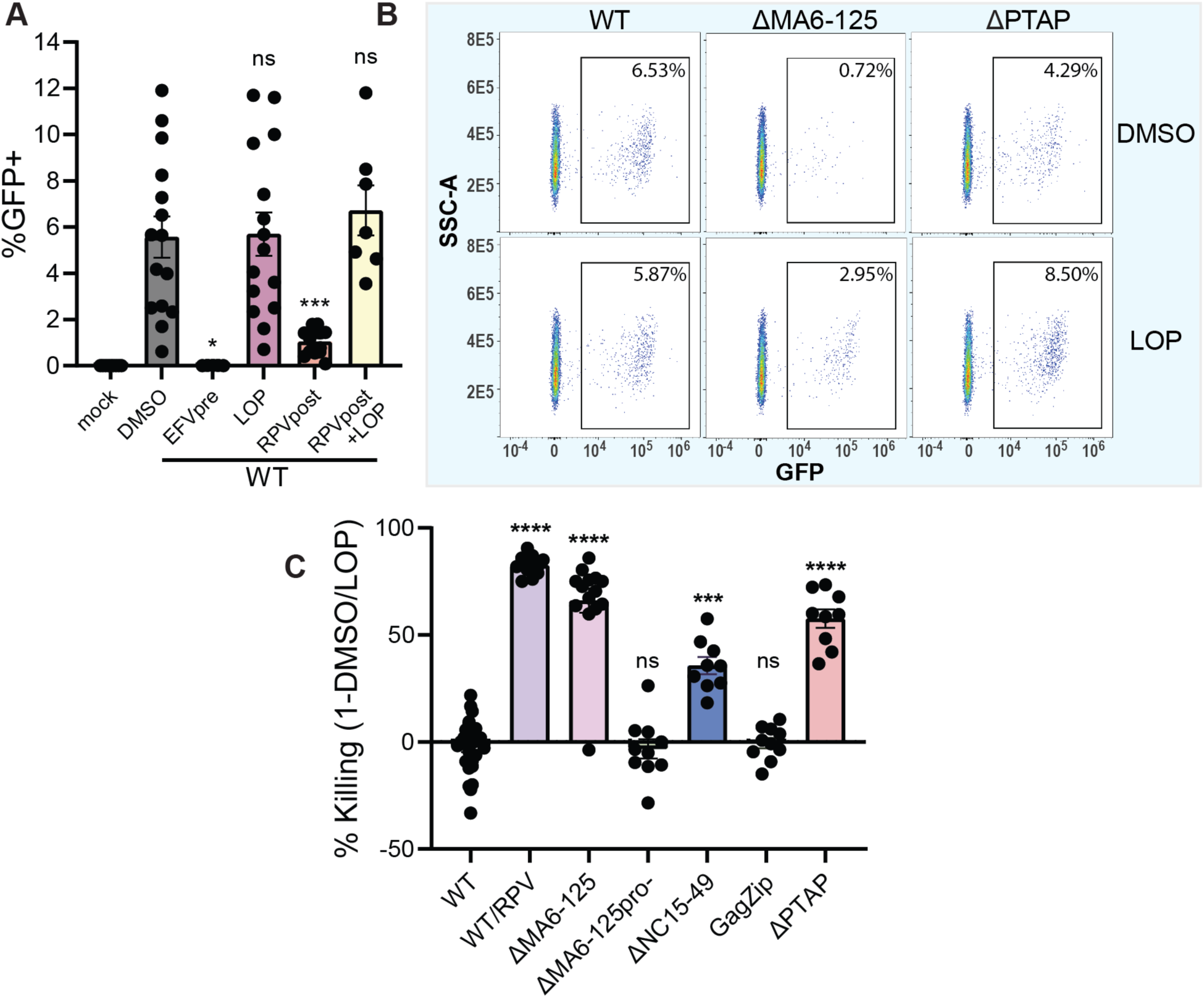
MA, NC, and p6 coordinate PR activity to evade CARD8 detection in CD4+ T cells. (**A**) Activated primary CD4+ T cells were infected with WT (LaiΔenvGFP/G) virus in the presence of 1 μM efavirenz (EFVpre), 1 μM lopinavir (LOP), and/or 5 μM rilpivirine (RPV, added 24h prior to harvest), with plot showing GFP positivity at 3 dpi. (**B**) Sample flow cytometry plots outlining how % killing was calculated from DMSO- and LOP-treated cultures. (**C**) Primary activated CD4+ T cells were infected with the indicated virus and PR-driven cytotoxicity was calculated by comparing GFP positivity in DMSO/LOP-treated cultures at 3 dpi (WT/RPV: WT virus infection treated with 5 μM RPV 24h pre-harvest compared to LOP+RPV). Data is shown as mean +/- SEM with each data point representing a different primary cell donor across over five independent experiments. Flow cytometry plots in (**B**) are from one representative primary cell donor. One-way ANOVA was used with Dunnett’s post-test with statistics referring to comparison with DMSO (**A**) or comparison with WT (**C**): ns: p≥0.05; *: p<0.05; ***: p<0.001; ****: p<0.0001.

To quantify PR-mediated cytotoxicity of Gag-mutant viruses, we calculated a % killing value comparing GFP positivity in DMSO-treated cultures to LOP-treated cultures (**Fig. 5B**). As expected, infection of activated CD4+ T cells with ΔMA6-125 or ΔPTAP viruses resulted in a significant increase in death of GFP+ infected cells relative to the LOP-treated control (**Fig. 5C**), similar in magnitude to that observed upon RPV treatment of WT infected CD4+ T cells. No death of GFP+ cells was observed if the ΔMA6-125 virus also carried the inactivating PR mutation, confirming that the cytotoxicity observed was not due to PR inhibitor treatment. Surprisingly, in contrast to myeloid cells, we now observed modest PR-mediated cell death upon infection with the ΔNC15-49 mutant virus. Intriguingly, this NC-modulated cell death was prevented in GagZip mutant virus infection, highlighting the important interplay between successful virus assembly and release and evasion of CARD8 sensing in CD4+ T cells. In total, these results indicate that primary CD4+ T cells are susceptible to CARD8-mediated pyroptotic cell death if virus assembly is dysregulated.

### Previously documented host-adaptive mutations in MA and p6 prevent pyroptosis of infected cells

Since structural mutants of Gag activate CARD8 via dysregulation of PR activation at the viral assembly site, it is likely that HIV-1 assembly is subject to innate immune pressure in human cells. Hence, we hypothesized that polymorphisms in Gag sequences might impact viral assembly and therefore modulate the ability of HIV-1 to evade CARD8 activation. For instance, near-universal adaptations in MA occurred during zoonotic transformation of SIVcpz to HIV-1, which presumably occurred concurrently with the encounter of CARD8 immune pressure in human cells.(*28*) While chimpanzees encode a functional CARD8 sensor, chimpanzee CARD8-N does not contain a cleavage site efficiently targeted by HIV-1 or SIVcpz PR,(*29*) and therefore Gag-mediated regulation of PR during viral assembly in chimpanzee cells may have less restrictive innate immune pressures compared to assembly in human cells. In contrast, human CARD8-N is efficiently cleaved by both HIV-1 and SIVcpz PR. We therefore hypothesized that adaptations in HIV-1 MA compared to SIVcpz may have arisen to combat innate immune pressure targeting virus assembly in human cells. To test this hypothesis, we employed a chimeric HIV-1 virus where HIV-1 MA sequence was replaced with SIVcpz MA (CPZ-MA) (**Fig. 6A**).(*16*) Intriguingly, when we tested PR-driven cytotoxicity of this chimaera in activated primary CD4+ T cells, we observed significantly greater PR-driven cytotoxicity (**Fig. 6B**). To probe this effect more closely at the single residue level, we mutated the 30K residue of Lai-MA to the chimpanzee-ancestral M (K30M), as this site is among the most universally conserved human-adaptive mutations that occurred during zoonosis, and M is the most common ancestral residue in SIVcpz.(*30*) Remarkably, we observed a significant increase in PR-induced cytotoxicity with K30M virus infection (**Fig. 6C**), implying that at least part of the function of this near-universal adaptation in HIV-1 MA is to prevent CARD8 activation during viral assembly in human cells.

**Fig. 6:**
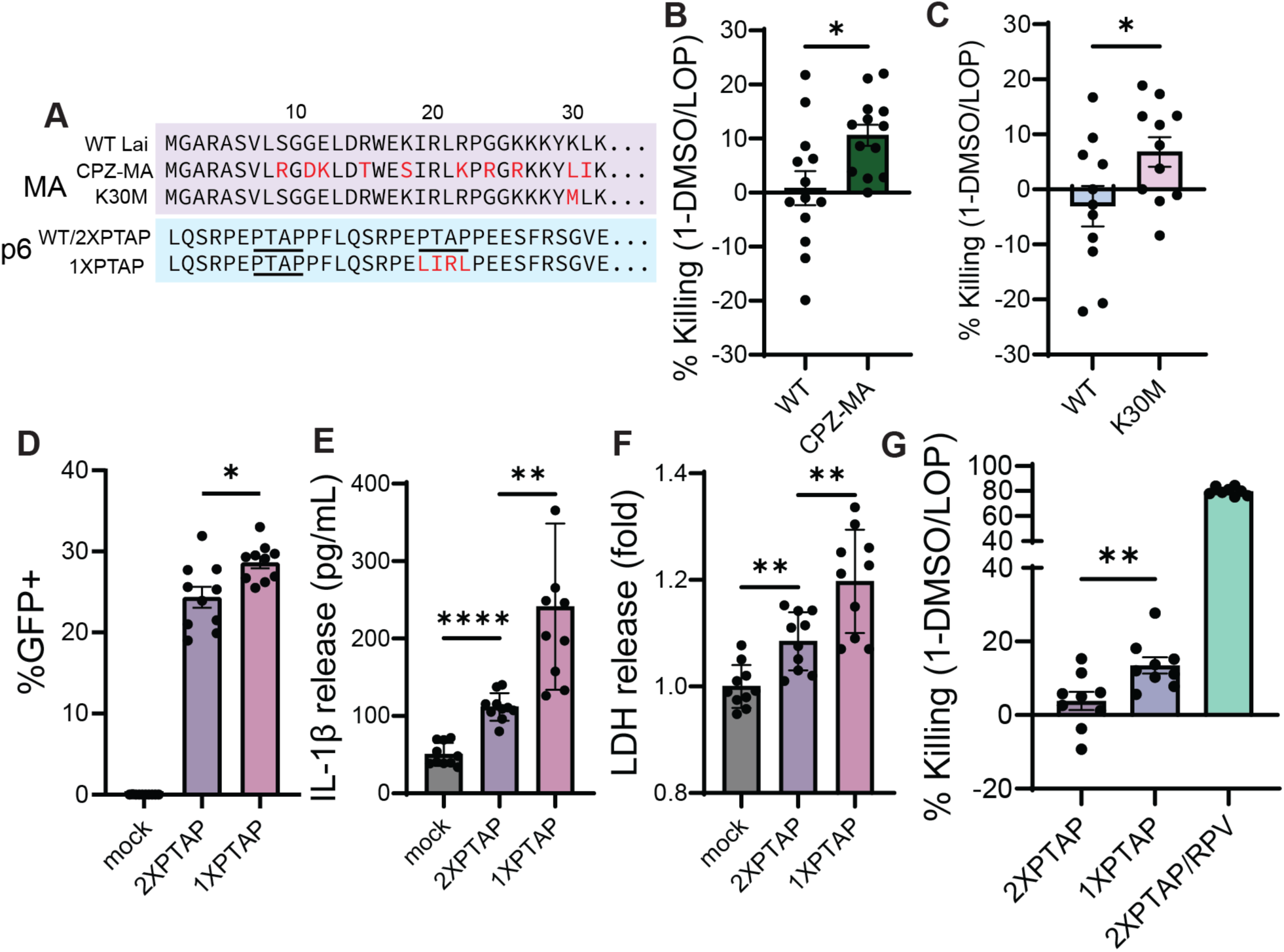
Human-adapted motifs in HIV-1 Gag reduce pyroptotic cell death during viral assembly. (**A**) Schematic outlining the MA and p6 mutant sequences used in these experiments compared to HIV-1 Lai (WT). Note that CPZ-MA has additional variations in MA not shown. WT and 2XPTAP refer to the same virus (LaiΔenvGFP/G). (**B**) Primary CD4+ T cell PR killing assay performed with WT and CPZ-MA virus or (**C**) with WT and K30M virus. Plots (**D**-**F**) show GFP positivity, IL-1β release, and LDH release in TNFα-treated PMA/THP cells co-infected with single-round HIV-1 and SIV3+ VLPs at 3 dpi. (**G**) Primary CD4+ T cell PR killing assay (2XPTAP/RPV: 2XPTAP virus infection treated with 5 μM RPV 24h pre-harvest). Data is shown as mean +/- SEM. (**B**/**C**/**G**) Data points are individual primary cell donors across over three independent experiments for each panel. (**D**-**F**) Data points are individual wells across three independent experiments. Data was analyzed with paired t-test (**B**) or one-way ANOVA with Dunnett’s post-test (**C**) or Tukey’s post-test (**D**-**G**): ns: p≥0.05; *: p<0.05; **: p<0.01; ****: p<0.0001.

We next turned our attention to human-adapted variants of HIV-1 p6. Multiple studies have shown that duplication of the Tsg101-recruiting PTAP motif in p6 confers faster virus budding and a replicative advantage in primary isolates of HIV-1.(*31–33*) Indeed, the lab-adapted Lai clone used in these studies contains two PTAP motifs, possibly attained after serial passage in human primary cell cultures vulnerable to CARD8 activation. Since disruption of both Lai PTAP motifs resulted in a dramatic enhancement of CARD8 activation during viral assembly (**Fig. 4**), we hypothesized that PTAP motif duplication could have possibly arisen to modulate CARD8 sensing during viral assembly. To address this question, we generated a single-PTAP mutant by mutating only one of the PTAP motifs in HIV-1/Lai to inert LIRL sequence (1XPTAP) and hypothesized that it would have a greater CARD8 activation phenotype compared to the wild-type virus (WT) containing two PTAP motifs (2XPTAP) (**Fig. 6A**). Indeed, when we infected PMA/THP macrophages with 1XPTAP virus, we observed IL-1β and LDH secretion levels greater than WT (2XPTAP) virus infection (**Fig. 6D**-**F**). Further, when we tested PR-induced cell death in primary activated CD4+ T cells, we found significantly increased PR-driven cell killing in 1XPTAP infection compared to WT (2XPTAP) virus infection (**Fig. 6G**). In addition, we analyzed the extent of PR-driven killing between the common HIV-1 laboratory isolates Lai (here called WT) and NL4-3, as one of the few striking differences between the Gag sequences of these two closely-related isolates is the duplication of the PTAP motif in Lai p6. Remarkably, consistent with our result with the 1XPTAP Lai mutant, we noted a significant enhancement in PR-driven killing in single-PTAP NL4-3 infection compared to WT Lai (2XPTAP) (**Fig. S3**). These results suggest that in addition to MA, the p6 region of Gag is also subject to selective pressure to improve survival of infected cells.

## DISCUSSION

In this report, we demonstrate that Gag-mediated virus assembly provides spatial and temporal control of PR activation until the final membrane scission step to prevent CARD8 sensing during productive HIV-1 infection. In the case of MA, we hypothesize a model that MA selects not only the PM but additionally selects specific lipid microdomains within the PM for proper initiation of Gag-pol auto-processing and viral assembly. Notably, we observed enhanced auto-processing of Gag-Pol and a concomitant increase in processed p66^RT^ in the cytoplasm of ΔMA6-125-transfected HEK293T cells, indicating abnormal initiation of PR activity. This model is supported by findings demonstrating that HIV-1 buds selectively from sphingolipid-enriched lipid microdomains (indicating that MA localizes viral assembly to specific lipid environments)(*34*) and also by findings indicating that inactivation of PM lipid-modifying enzymes or mutations in the membrane-interacting HBR have drastic effects on Gag processing,(*35–37*) implying a crosstalk between MA-engaged lipids and PR activity. We speculate that membrane fluidity determined by local lipid microenvironment is altered during virus assembly in the absence MA HBR or in the presence of HBR mutations, which might increase the probability of Gag-Pol monomer interactions resulting in enhanced propensity for dimerization and initiation of PR activity and maturation.

In contrast, PTAP-deficient viral assembly proceeds past the point where active PR is generated, but is halted before the final budding step which evidently must complete within a short timeframe to avoid escape of catalytically-active PR from the assembly site, which triggers CARD8 activation. This finding is consistent with previous reports relating mutations in p6 to a reduced retention of viral enzymes during assembly.(*24, 25*) Separately, it is intriguing that NC has a divergent phenotype in macrophages and CD4+ T cells in terms of CARD8 evasion. However, HIV-1 NC has been previously reported to contain an ALIX-binding motif,(*22*) and virus budding during NC-deficient assembly has been shown to be slower relative to WT via an unknown mechanism dependent on PR activation, implicating NC as a regulator of PR activity and budding.(*38*) Indeed, NC’s role in CARD8 evasion in T cells specifically might reflect differential usage of Tsg101- and ALIX-driven budding pathways between myeloid and lymphoid cells infected with HIV-1.(*39*)

To become pandemic in the human population, HIV-1 had to simultaneously antagonize multiple mechanisms of restriction between great apes and humans. Perhaps unsurprisingly, then, most zoonotic transmission events of SIVcpz or SIVgor from great apes to humans did not result in the establishment of widely successful HIV-1 lineages. This work reveals that, in addition to the previously described species-specific restrictions such as tetherin and the APOBECs, CARD8 activation by assembly site-generated PR is an additional layer of restriction between chimpanzees and humans. Recent research has shown that chimpanzee CARD8 is not sensitive to lentiviral PR cleavage,(*29*) implying that CARD8 evasion was an innate immune barrier encountered by zoonotic SIVcpz immediately at the cross-species transmission event to humans. It is worth noting that our CPZ-MA chimaera is based on the SIVcpz isolate TAN2, which was sampled from *Pan troglodytes schweinfurthii* (*P.t.s.*) in Tanzania, not from *Pan troglodytes troglodytes* (*P.t.t.*) in central Africa which are believed to be the hosts of the direct ancestors of HIV-1.(*40*) The presence of innate immune-activating motifs in *P.t.s.* SIVcpz viruses could partially explain why only viruses descended from central African *P.t.t.* achieved pandemicity in the human population.

This work also has important implications for drug discovery and therapeutic approaches to tame the HIV-1 reservoir in ART-suppressed people with HIV. NNRTIs have recently garnered interest for drug-induced killing of HIV-infected cells via CARD8 activation. Our results show that MA and p6 could be attractive targets to either enhance NNRTI-driven cell killing or induce CARD8 activation themselves. Therefore, drug development strategies should consider modulation of virus assembly to achieve maximum CARD8-mediated elimination of infected cells.

## MATERIALS AND METHODS

### Viruses, plasmids, and cloning

To make a pro-IL-1β expression vector, the pro-IL-1β coding sequence was excised from pLV-mTurqoise2-IL1β-mNeonGreen plasmid (Addgene cat. # 166783) via *Xho*I/*Bam*HI restriction sites and introduced into pcDNA3.1. The pCI-Caspase1 expression plasmid was also received from Addgene (cat. # 41552). For transfection experiments, the ‘empty’ vector transfected was a pCS2 plasmid encoding eYFP. The single-round proviral molecular clones LaiΔenvGFP (here called ‘WT’) and the matrix mutant LaiΔMA6-125ΔenvGFP with inactivating deletions in *env* and GFP in place of *nef* have been described previously.(*16*) NL4-3ΔenvGFP has been described previously (received from NIH/NIAID HIV Reagent Program, cat. # ARP-11100). The M10 and G2A mutations on the Lai backbone were previously reported.(*16*) The Lai pro- (in which residues 25, 49, and 50 are mutated from D/G/I to K/W/W) and Lai VK181/2AA (CA-CTD dimer interface mutant) clones were generous gifts from Dr. Jaisri Lingappa at the University of Washington.(*16*) The pro-mutation was swapped into the LaiΔMA6-125ΔenvGFP plasmid via *Apa*I/*Sal*I. The VK181/2AA CA mutation was swapped into the LaiΔMA6-125ΔenvGFP plasmid via *Age*I/*Apa*I. To make a CARD8-targeting shRNA lentivector, CARD8 shRNA was excised from a construct purchased from Millipore Sigma (cat. # TRCN0000118329, target sequence GCACAAACAGATACAGCGTTT) and introduced into a hygromycin-resistance pLKO.1 vector via *Spe*I/*Mfe*I. To generate G2A-ΔMA6-125, ΔNC15-49, and ΔPTAP, a Gibson-based strategy was used. First, two fragments were generated from LaiΔenvGFP using PCR (employing Phusion High-Fidelity DNA polymerase, New England Biolabs, cat. # M0530) with primers #1-12 included in **Table S1**. N and C terminal fragments of the correct size were gel-purified. Then, Gibson assembly was performed using the two amplified fragments (Gibson Assembly Cloning Kit, New England Biolabs, cat. # E5510S). 1 μL of the Gibson assembly reaction was then used as template in a final PCR reaction using the furthest 5’ and 3’ primers, with final fragments of the correct size gel-purified, restriction digested, and ligated into LaiΔenvGFP (G2A/ΔMA6-125: *BssH*II/*Age*I, ΔNC15-49: *Age*I/*Pas*I, ΔPTAP: *Apa*I/*Pas*I). To inactivate PTAP motifs, PTAP (CCAACAGCCCCA) was mutated to the inert sequence LIRL (CTAATACGACTA). ΔNC15-49 was generated by deleting residues 15-49 of NC (inclusive), from the beginning of the first zinc finger to the end of the second. Lai-GagZip was a generous gift from Dr. Jaisri Lingappa, which was ligated into LaiΔenvGFP via *Age*I/*Sal*I.(*23*) To make the K30M mutation in LaiΔenvGFP, the QuikChange II Site-Directed Mutagenesis Kit (Agilent Technologies, cat. # 200523) was used employing primers in **Table S1**. To make LaiΔenvGFP/1XPTAP, a DNA fragment was purchased from Genewiz-Azenta with the second PTAP in Lai-p6 mutated to LIRL, which was digested and ligated into LaiΔenvGFP via *Apa*I/*Bcl*I. The CPZ-MA plasmid was described previously.(*16*) All plasmid sequences were confirmed via Sanger sequencing (Genewiz-Azenta) and/or restriction digestion prior to use.

All viruses were produced via calcium phosphate-mediated double/triple transfection of HEK293T cells in dishes followed by ultracentrifugation over a 20% sucrose cushion, extensively described elsewhere.(*16*) To generate single-round VSV-G-pseudotyped viruses, all mutant viral plasmids (except WT) were co-transfected with an HIV-1 packaging plasmid (psPAX2) and a VSV-G-encoding expression vector. Lentivectors for cell line generation were produced via triple transfection. Vpx-containing SIV3+ particles were obtained by double transfection of SIV3+ plasmid and VSV-G, which were titered for capsid content using p27 ELISA (XpressBio, cat. # SK845) before use. The VSV-G, SIV3+, and psPAX2 plasmids have been described before.(*16, 18*)

### Cell lines

HEK293T cells and THP-1 cells were obtained from American Type Culture Collection (ATCC). TZM-bl cells used for titering viruses were obtained from the NIH/NIAID HIV Reference Reagent Program (cat. # ARP-8129). The HEK293T-CARD8KO CRISPR-knockout cell line was a generous gift from Dr. Liang Shan at Washington University in St. Louis. HEK293T and TZM-bl cell lines were maintained in DMEM high glucose medium (Gibco, cat. # 11965) plus 10% fetal bovine serum (Gibco, cat. # A5670) and 1% Pen-Strep (Gibco, cat. # 15140) (D10 medium), while THP-1 cells were maintained in RPMI1640 (Gibco, cat. # 11875) plus 10% FBS and Pen-Strep (R10 medium). To generate knockdown THP-1 cell lines, 1E6 THP-1 cells were infected with 400 ng p24^Gag^ lentivector encoding shScramble or shCARD8 via spinoculation in polybrene-containing medium. Transformed cells were selected and maintained in medium containing 400 μg/mL hygromycin (Invitrogen, cat. # 10-687-010). All cell lines used in these studies were routinely tested for mycoplasma contamination.

### Isolation and differentiation of primary cells

Peripheral blood mononuclear cells (PBMCs) were isolated from de-identified leukopacks obtained from NY Biologics as described previously.(*16*) Primary CD4+ T cells were obtained either from whole PBMCs or CD14-flowthrough and positively isolated using anti-CD4 magnetic beads (Miltenyi Biotech, cat. # 130-045-101). CD4+ T cells were stimulated in R10 medium with 2% phytohemagglutinin (PHA, Gibco, cat. # 10576015) and 50 U/mL IL-2 (NIH/NIAID HIV Reference Reagent Program, cat. # 136) for two days before washing in DPBS and resuspension in R10 medium with 50 U/mL IL-2.

### Infections

THP-1 cells in suspension were differentiated into PMA/THP-1 macrophage-like cells by treatment with phorbol 12-myristate 13-acetate (PMA) (100 nM, Sigma-Aldrich, cat. # P8139) for 2 days before infection. PMA/THP-1 cells were seeded at 2.5E5 cells/well in 24-well plates for infection and co-infected with HIV-1 and 5 ng p27 Vpx-containing SIV3+ VLPs in the presence of polybrene (10 µg/mL, Millipore-Sigma, cat. # TR1003G) by spinoculation at 2300 rpm for 1 hour at room temperature before incubation for 2-3 hours at 37°C. After the incubation, cells were washed with DPBS to remove unbound viral particles and 500 µL fresh R10 media was added supplemented with 10 ng/mL TNFα (Peprotech, cat. # 300-01A). Cells were harvested at 3 dpi. Supernatants were collected and clarified by centrifugation (1200 rpm, 5 min) before analysis and cells were harvested for RIPA lysis or flow cytometry.

Activated primary CD4+ T cells were infected in the presence of polybrene in 0.5 mL of R10 media in 24-well plates at 2E6 cells/well. As with macrophages, spinoculation was performed for 1 hour at room temperature before incubation for 2-3 hours at 37°C. Infected cells were then thoroughly washed with DPBS, before cultures were split in two and 1E6 cells were cultured in 300 µL DMSO- or LOP-containing R10/IL-2 medium in flat-bottom 96-well plates. T cells were harvested for flow cytometry analysis at 3 dpi. % killing in primary CD4+ T cell experiments was calculated using the formula:

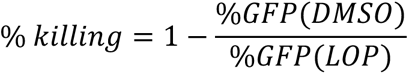

All infections were performed at MOI 1 based on TZM-bl titer.

### Antibodies and drugs

Antibodies used for Western blot analysis are as follows: IL-1β (mouse clone # 2805, R&D Systems, cat. # MAB601, used at 1:1000), p24^Gag^ (mouse clone # p24-2, NIH HIV Reagent Program, cat. # 6457, used at 1:1000), p24^Gag^ (rabbit polyclonal, Immuno Diagnostics, cat. # 1303), human HIV-Ig (human polyclonal, NIH, cat. # ARP-3957, used at 1:2000), β-actin (mouse clone # AC-15, Invitrogen, cat. # AM4302, used at 1:5000), β-actin (rabbit polyclonal, Sigma-Aldrich, cat. # A2066, used at 1:5000), CARD8-N (rabbit poyclonal, abcam, cat. # ab194585, used at 1:1000), CARD8-C (rabbit polyclonal, abcam, cat. # ab24186, used at 1:1000). Secondary antibodies used were goat anti-mouse DyLight 680 (Invitrogen, cat. # 35518, used at 1:10,000), goat anti-rabbit DyLight 800 (Invitrogen, cat. # SA5-35571, used at 1:10,000), and goat anti-human DyLight 800 (Rockland, cat. # 609-145-123, used at 1:10,000).

Antiretrovirals used include efavirenz (NIH/NIAID HIV Reference Reagent Program, cat. # 4624), lopinavir (NIH/NIAID HIV Reagent Program, cat. # HRP-9481), saquinavir (NIH/NIAID HIV Reagent Program, cat. # ARP-4658), rilpivirine (NIH/NIAID HIV Reagent Program, cat. # HRP-12147), tenofovir (Medkoo Biosciences, cat. # 318800). Antiretrovirals were used at 1 μM, except for tenofovir used at 30 μM, or rilpivirine/efavirenz used at 5 μM to induce CARD8-mediated cell death. Rilpivirine/efavirenz added at 5 μM was added at 24 hours pre-harvest. Ilaprazole was used at 1 or 10 μM (Selleck Chemicals, cat. # S3666). Valboro-Pro (VbP, InvivoGen, cat. # tlrl-vbp-10) was used at 10 μM.

### HEK293T transfection

To analyze IL-1β cleavage and secretion *in situ*, HEK293T cells were transfected using *Trans*IT-293 transfection reagent (Mirus Bio, cat. # MIR 2704). 2.5E5 HEK293T cells per well were seeded in 1 mL of D10 media in 12-well plates. The next day, cells were transfected according to the manufacturer’s instructions. Each well was transfected with 5 ng of plasmid encoding CASP1, 200 ng of plasmid encoding pro-IL-1β, and 200 ng of control plasmid or HIV-1 construct. Cells were harvested roughly 24 hours post-transfection. Supernatants were collected and clarified by centrifugation. Cell lysates were harvested for Western blot analysis by washing with DPBS before lysis in RIPA buffer. To transfect HEK293T cells for virus assembly analysis, one 10-cm dish was transfected with 10 µg of Δenv proviral plasmid alone. Cells were lysed 48 hours post-transfection and viruses were pelleted in an ultracentrifuge using sucrose as described above before lysis in RIPA buffer (150 μL).

### ELISA

Human IL-1β in clarified culture supernatants was quantified using the IL-1β DuoSet ELISA kit (R&D Systems, cat. # DY201) according to the manufacturer’s protocol after inactivation in DPBS buffer with 10% normal calf serum (Invitrogen, cat. # 26170043) and 0.5% TX-100 detergent. HIV-1 p24^Gag^ levels were quantified using in-house ELISA as previously described after a similar inactivation step.(*16*)

### LDH release assay

Lactate dehydrogenase (LDH) release into culture supernatants was quantified using the CytoTox 96 Non-Radioactive Cytotoxicity Assay (Promega, cat. # PR-G1780) according to the manufacturer’s instructions after inactivation in DPBS with 0.5% TX-100. LDH assays were stopped with Stop Solution after 15-30 minutes of development and read at 490 nm.

### Western blot analysis

To perform quantitative Western blot analysis, 30 μg of protein (or 10 μL of virus lysate) was normalized to the same volume in DPBS/loading buffer before boiling and loading into Mini-PROTEAN TGX Precast Protein Gels (BioRad, 10% - cat. # 4561034, or 4-20% gradient – cat. # 4561094) along with a protein ladder (PageRuler Prestained NIR Protein Ladder, Thermo Scientific, cat. # 26635). To resolve small proteins for analysis of p25/p24^Gag^ levels, a 12.5% acrylamide hand-poured gel was used. Blots were blocked in LI-COR Intercept blocking buffer (LI-COR, cat. # NC1660556) before membrane staining and imaging on a LI-COR Odyssey CLx imager.

### Flow cytometry

Cells to be analyzed by flow cytometry were first washed with DPBS before fixing with 4% paraformaldehyde (PFA) for 15-30 minutes. PFA was washed out and cells were analyzed on a Cytek Aurora spectral analyzer. GFP was measured in the B2 channel after gating on single cells via FSC and SSC. Data was analyzed and plots were created in FlowJo software.

### Alignment and phylogenetic analysis

Amino acid alignments were generated using Clustal Omega (European Bioinformatics Institute).(*41*)

### Statistics

Unless otherwise specified, statistical significance was assessed in GraphPad Prism 10 using one-way ANOVA with Tukey’s or Dunnett’s post-test. For experiments with comparative cell lines (i.e. shScramble versus shCARD8), two-way ANOVA was used. For comparisons of only two groups, unpaired or paired T-test was used. Plots show mean of the data points ± SEM. Data points for primary cell experiments represent individual donors across multiple experiments, while data points in cell line experiments represent multiple wells in the same experiment, or individual wells across multiple experiments. ns: p≥0.05; *: p<0.05; **: p<0.01; ***: p<0.001; ****: p<0.0001.

## Acknowledgements

The authors thank the BU-CAMED Flow Cytometry Core Facility for technical assistance, the Shan laboratory at Washington University in St. Louis for their generous gift of HEK293T-CARD8KO cells, and the NIH/NIAID HIV Reagent Program and associated investigators for reagents used in this study. This work was supported by NIH grants R01DA059952 (SG), R01AI187175 (SG), R01DA055488 (SG), R01DA051889 (SG), and 5P30AI042853 (SG and HA). The authors declare that they have no competing interests.

## Author Contributions

I.H., H.A., and S.G. designed the experiments. I.H. and S.G. wrote the manuscript. I.H. performed the experiments with support from J.H., A.Q.M., and H.A.

## Data Availability

All data needed to evaluate the conclusions in the paper are present in the paper and/or the Supplementary Materials. The authors declare that the data that support the findings of this study are available from the corresponding author upon reasonable request.

## SUPPLEMENTARY FIGURE LEGENDS

**Fig. S1:**
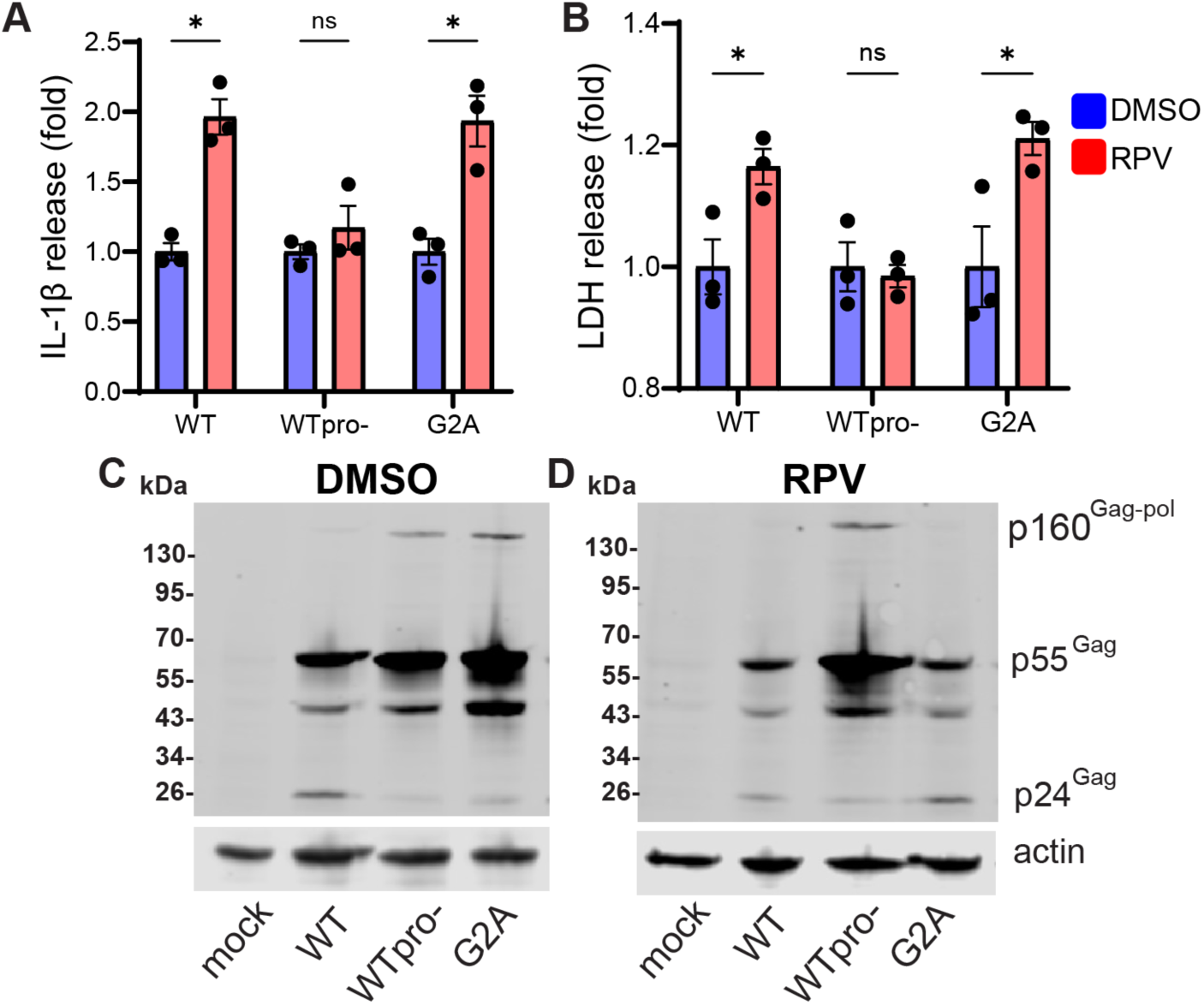
CARD8 can sense NNRTI-activated HIV-1 PR in the cytosol or at the membrane assembly site. (**A** and **B**) IL-1β and LDH release (normalized to DMSO condition) of TNFα-treated PMA/THP cells co-infected with single-round VSV-G-pseudotyped HIV-1Δenv/GFP and SIV3+ VLPs, stimulated with DMSO or 5 μM RPV 24h prior to harvest (3 dpi total). Means +/- SEM are shown and statistics refer to multiple unpaired t-tests from one experiment. (**C**/**D**) Western blots show cell lysates of PMA/THP cells in **A**/**B**, harvested at time of supernatant collection and probed with anti-p24^Gag^ antibody.

**Fig. S2:**
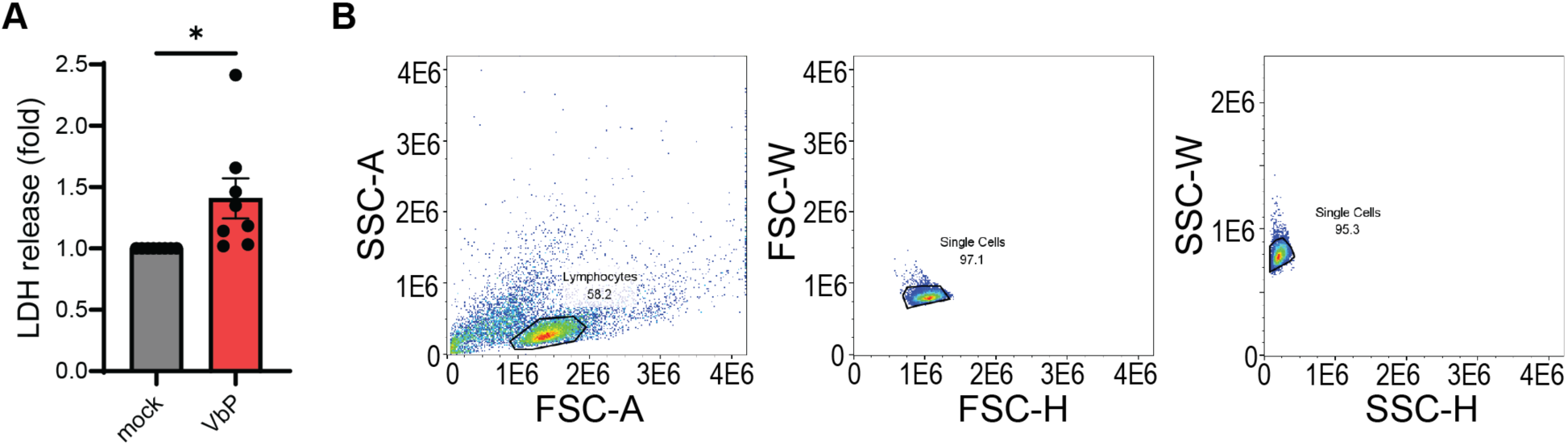
VbP-induced pyroptosis in CD4+ T cells. (**A**) LDH secretion from activated CD4+ T cells used for cytotoxicity experiments 24h post-10 μM VbP treatment, with mean +/- SEM shown and paired t-test comparison of individual primary cell donors across three independent experiments. (**B**) Sample gating strategy for assessing GFP positivity in single CD4+ T cells.

**Fig. S3:**
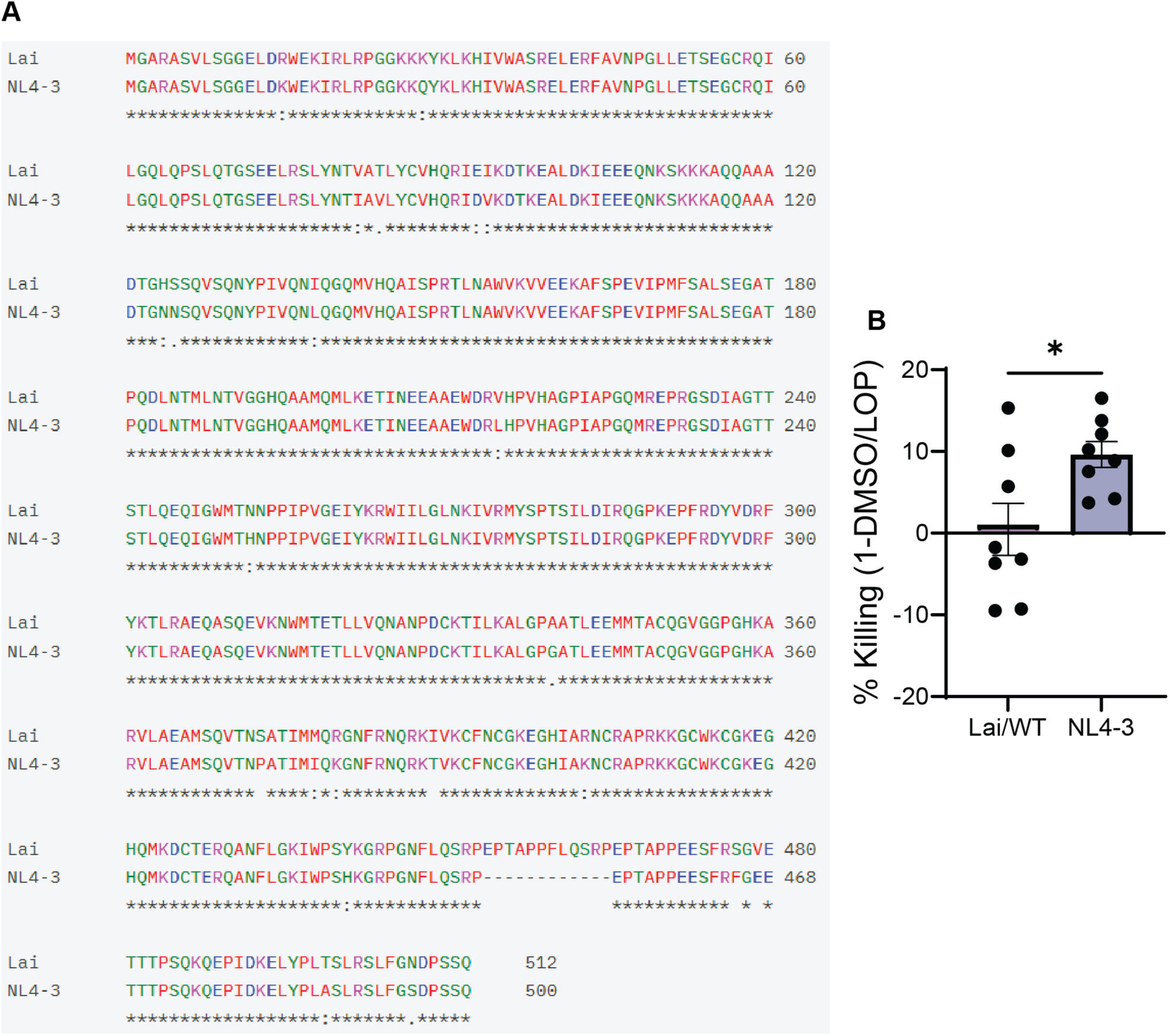
Differential CARD8 evasion between HIV-1 Lai and NL4-3. (**A**) Clustal alignment of Lai (WT) and NL4-3 Gag sequences. (**B**) CD4+ T cell PR killing assay with cells infected with single-round Lai (WT) or single-round NL4-3 GFP-encoding viruses. Mean +/- SEM is shown. Data points are individual primary cell donors with paired t-test comparison across three independent experiment.

**Table S1:**
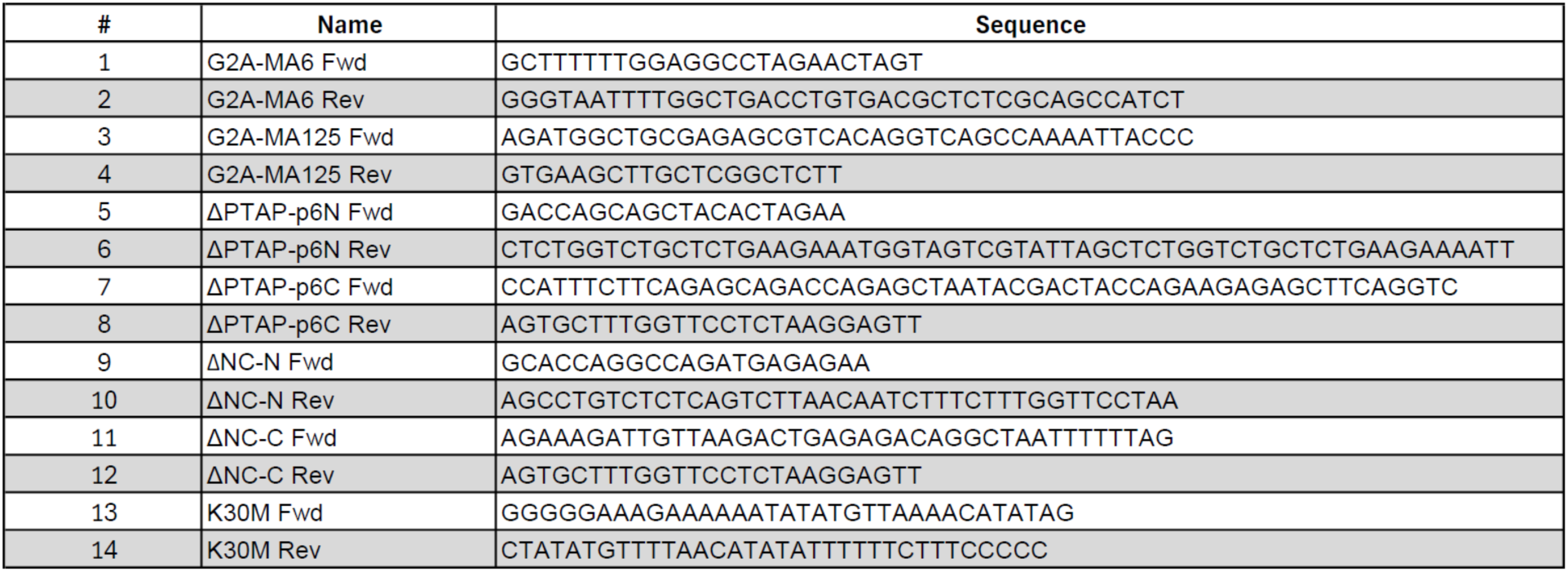
Primers used for cloning.

