## Supplementary file for "Evasion of CARD8 Activation During HIV-1 Assembly"

### SUPPLEMENTARY FIGURE LEGENDS

**Figure S1**

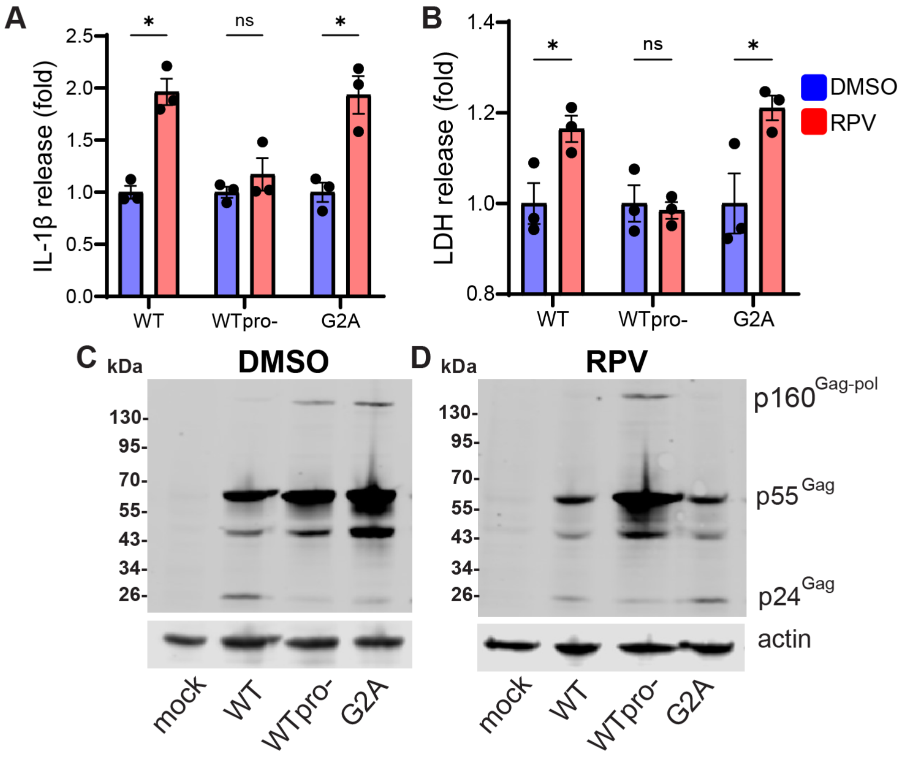


**Fig. S1: CARD8 can sense NNRTI-activated HIV-1 PR in the cytosol or at the membrane assembly site.** (**A** and **B**) IL-1β and LDH release (normalized to DMSO condition) of TNFα-treated PMA/THP cells co-infected with single-round VSV-G-pseudotyped HIV-1Δenv/GFP and SIV3+ VLPs, stimulated with DMSO or 5 µM RPV 24h prior to harvest (3 dpi total). Means +/- SEM are shown and statistics refer to multiple unpaired t-tests from one experiment. (**C**/**D**) Western blots show cell lysates of PMA/THP cells in **A**/**B**, harvested at time of supernatant collection and probed with anti-p24$^{Gag}$ antibody.

**Figure S2**

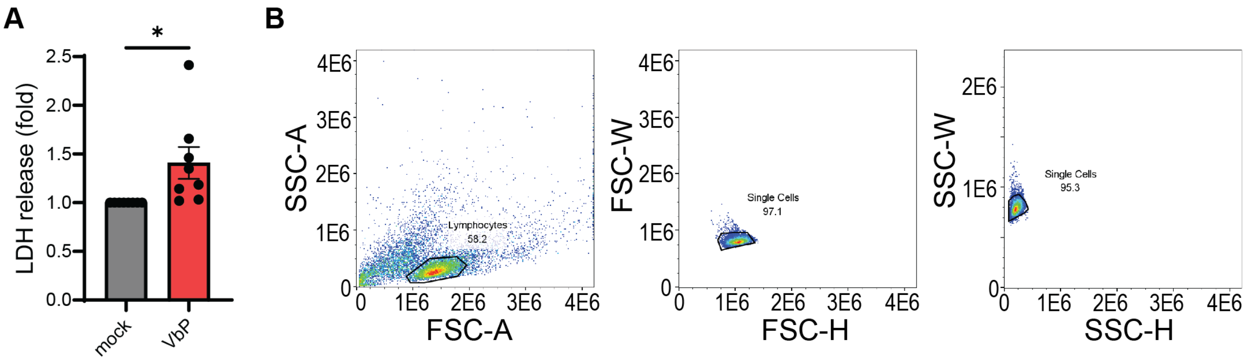


**Fig. S2: VbP-induced pyroptosis in CD4+ T cells.** (**A**) LDH secretion from activated CD4+ T cells used for cytotoxicity experiments 24h post-10 μM VbP treatment, with mean +/- SEM shown and paired t-test comparison of individual primary cell donors across three independent experiments. (**B**) Sample gating strategy for assessing GFP positivity in single CD4+ T cells.

**Figure S3**

**A**

```
Lai         MGARASVLSGGELDRWEKIRLRPGGKKKYKLKHIVWASRELERFAVNPGLLETSEGCRQI 60
NL4-3       MGARASVLSGGELDKWEKIRLRPGGKKQYKLKHIVWASRELERFAVNPGLLETSEGCRQI 60
            **************:************:********************************

Lai         LGQLQPSLQTGSEELRSLYNTVATLYCVHQRIEIKDTKEALDKIEEEQNKSKKKAQQAAA 120
NL4-3       LGQLQPSLQTGSEELRSLYNTIAVLYCVHQRIDVKDTKEALDKIEEEQNKSKKKAQQAAA 120
            *********************:*.*********::*************************

Lai         DTGHSSQVSQNYPIVQNIQGQMVHQAISPRTLNAWVKVVEEKAFSPEVIPMFSALSEGAT 180
NL4-3       DTGNNSQVSQNYPIVQNLQGQMVHQAISPRTLNAWVKVVEEKAFSPEVIPMFSALSEGAT 180
            ***:.************:******************************************

Lai         PQDLNTMLNTVGGHQAAMQMLKETINEEAAEWDRVHPVHAGPIAPGQMREPRGSDIAGTT 240
NL4-3       PQDLNTMLNTVGGHQAAMQMLKETINEEAAEWDRLHPVHAGPIAPGQMREPRGSDIAGTT 240
            **********************************:*************************

Lai         STLQEQIGWMTNNPPIPVGEIYKRWIILGLNKIVRMYSPTSILDIRQGPKEPFRDYVDRF 300
NL4-3       STLQEQIGWMTHNPPIPVGEIYKRWIILGLNKIVRMYSPTSILDIRQGPKEPFRDYVDRF 300
            ***********:************************************************

Lai         YKTLRAEQASQEVKNWMTETLLVQNANPDCKTILKALGPAATLEEMMTACQGVGGPGHKA 360
NL4-3       YKTLRAEQASQEVKNWMTETLLVQNANPDCKTILKALGPGATLEEMMTACQGVGGPGHKA 360
            ***************************************.********************

Lai         RVLAEAMSQVTNSATIMMQRGNFRNQRKIVKCFNCGKEGHIARNCRAPRKKGCWKCGKEG 420
NL4-3       RVLAEAMSQVTNPATIMIQKGNFRNQRKTVKCFNCGKEGHIAKNCRAPRKKGCWKCGKEG 420
            ************ ****:*:******** *************:*****************

Lai         HQMKDCTERQANFLGKIWPSYKGRPGNFLQSRPEPTAPPFLQSRPEPTAPPEESFRSGVE 480
NL4-3       HQMKDCTERQANFLGKIWPSHKGRPGNFLQSRP------------EPTAPPEESFRFGEE 468
            ********************:************            *********** * *

Lai         TTTPSQKQEPIDKELYPLTSLRSLFGNDPSSQ     512
NL4-3       TTTPSQKQEPIDKELYPLASLRSLFGSDPSSQ     500
            ******************:*******.*****
```

**B**

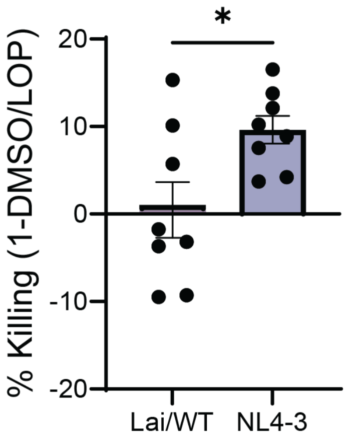


**Fig. S3: Differential CARD8 evasion between HIV-1 Lai and NL4-3.** (**A**) Clustal alignment of Lai (WT) and NL4-3 Gag sequences. (**B**) CD4+ T cell PR killing assay with cells infected with single-round Lai (WT) or single-round NL4-3 GFP-encoding viruses. Mean +/- SEM is shown. Data points are individual primary cell donors with paired t-test comparison across three independent experiments.

#### Table S1

| # | Name | Sequence |
|---|---|---|
| 1 | G2A-MA6 Fwd | GCTTTTTTGGAGGCCTAGAACTAGT |
| 2 | G2A-MA6 Rev | GGGTAATTTTGGCTGACCTGTGACGCTCTCGCAGCCATCT |
| 3 | G2A-MA125 Fwd | AGATGGCTGCGAGAGCGTCACAGGTCAGCCAAAATTACCC |
| 4 | G2A-MA125 Rev | GTGAAGCTTGCTCGGCTCTT |
| 5 | ΔPTAP-p6N Fwd | GACCAGCAGCTACACTAGAA |
| 6 | ΔPTAP-p6N Rev | CTCTGGTCTGCTCTGAAGAAATGGTAGTCGTATTAGCTCTGGTCTGCTCTGAAGAAAATT |
| 7 | ΔPTAP-p6C Fwd | CCATTTCTTCAGAGCAGACCAGAGCTAATACGACTACCAGAAGAGAGCTTCAGGTC |
| 8 | ΔPTAP-p6C Rev | AGTGCTTTGGTTCCTCTAAGGAGTT |
| 9 | ΔNC-N Fwd | GCACCAGGCCAGATGAGAGAA |
| 10 | ΔNC-N Rev | AGCCTGTCTCTCAGTCTTAACAATCTTTCTTTGGTTCCTAA |
| 11 | ΔNC-C Fwd | AGAAAGATTGTTAAGACTGAGAGACAGGCTAATTTTTTAG |
| 12 | ΔNC-C Rev | AGTGCTTTGGTTCCTCTAAGGAGTT |
| 13 | K30M Fwd | GGGGGAAAGAAAAAATATATGTTAAAACATATAG |
| 14 | K30M Rev | CTATATGTTTTAACATATATTTTTTCTTTCCCCC |

**Table S1:** Primers used for cloning.
